# Introducing Molecular Hypernetworks for Discovery in Multidimensional Metabolomics Data

**DOI:** 10.1101/2023.09.29.560191

**Authors:** Sean M Colby, Madelyn R Shapiro, Andy Lin, Aivett Bilbao, Corey D Broeckling, Emilie Purvine, Cliff A Joslyn

## Abstract

Orthogonal separations of data from high-resolution mass spectrometry can provide insight into sample composition and help address the challenge of complete annotation of molecules in untargeted metabolomics. “Molecular networks” (MNs), as used, for example, in the Global Natural Products Social Molecular Networking platform, are an increasingly popular computational strategy for exploring and visualizing molecular relationships and improving annotation. MNs use graph representations to show the relationships between measured multidimensional data features. MNs also show promise for using network science algorithms to automatically identify targets for annotation candidates and to dereplicate features associated to a single molecular identity. However, more advanced methods may better represent the complexity present in samples. Our work aims to increase confidence in annotation propagation by extending molecular network methods to include “molecular hypernetworks” (MHNs), able to natively represent multiway relationships among observations supporting both human and analytical processing. In this paper we first introduce MHNs illustrated with simple examples, and demonstrate how to build them from liquid chromatography-and ion mobility spectrometry-separated MS data. We then describe a method to construct MHNs directly from existing MNs as their “clique reconstructions”, demonstrating their utility by comparing examples of previously published graph-based MNs to their respective MHNs.

## Introduction

Small molecules play a crucial role in various biological processes and are widely studied in myriad fields such as drug discovery, environmental science, and human health explorations. However, the identification of small molecules in complex samples can be challenging due to the sheer number of potential detectable molecular configurations, measured by representative signatures, each often with high degrees of intersection. This challenge is further compounded by the presence of numerous interfering substances and the low concentrations of the target molecules.

To overcome these challenges, various measurement technologies have been developed to analyze small molecules in complex samples. The most widely used include liquid chromatography (LC), which separates molecules based on their physical and chemical properties; mass spectrometry and tandem mass spectrometry (MS/MS), which fragments molecules into characteristic pieces and measures the masses of the parent ion and corresponding fragments. Less commonly, though with burgeoning utility, ion mobility spectrometry (IMS) can be used to differentiate among compounds based on their gas-phase ion mobility, offering separation among molecules indistinguishable by LC, MS, or MS/MS, such as stereoisomers. Other techniques, such as nuclear magnetic resonance (NMR), infrared (IR), and ultraviolet (UV) spectroscopy, have the potential to further increase discriminatory power through consideration of additional dimensions. These developments have seen the rise of so-called “hyphenated” mass spectrometry analyses such as LC-MS/MS, LC-IMS-MS/MS, and IMS-IR-MS. These technologies generate data sets that are multidimensional, in the sense that multiple observational dimensions are generated for each feature, e.g. two for LC-MS/MS, or three for LC-IMS-MS/MS. Such multidimensional data can be challenging to analyze due to the large amount of information contained within. Additionally, the complexity of the samples can lead to difficulties in accurately identifying and quantifying the small molecules present.

Molecular networks (MNs) have been demonstrated as a useful tool for identifying small molecules in hyphenated MS data.^1–3^ MNs use mathematical graph structures (edges linking nodes) to represent the relationships among different compounds, including biochemical reactions, mass spectrometry features, structural similarities, or metabolite correlations. ^4^ Some of the most prominent MNs are used in the Global Natural Products Social Molecular Networking (GNPS)^3^ environment.^∗^ GNPS is an increasingly popular computational strategy for visualizing and interpreting interrelatedness of MS/MS spectra.

MNs offer an overview of the complex data: the network can highlight clusters of compounds that are likely to be related, which can narrow the search for molecules of interest. Additionally, the network can represent the relationships between different compounds, which can provide insight into the metabolism and biotransformation of the small molecules under study. MNs can also facilitate the integration of multiple types of data. For example, “feature-based molecular networks” (FBMNs) ^1^ incorporate joint information from both the LC and MS dimensions of the data, which can provide a more nuanced view of the small molecules present. This form of characterization can be particularly useful when working with complex samples, as it can help to differentiate between compounds that may be difficult to distinguish based on a single type of data.

Where MNs using mathematical graphs can provide significant value for representing complex data relationships, they exist as one method within a broader world of “network science” or “relational data” methods. This has at least two consequences:

- First, in relational data analysis, when considering how to approach multiple data dimensions (in our case, data dimensions measuring MS, LC, IMS, MS/MS, etc.), there can be considerable complexity in deciding which dimensions, and in particular which *combinations* of dimensions, should be attended to as either the objects to be focused on, or the relationships between those objects. These choices determine which particular network “view” of a relational data set is adopted. For example, the view of GNPS MNs is to attend to MS values as objects with similarities between their MS/MS spectra used to establish relationships; the views of GNPS FBMNs attend to joint LC-MS values as objects, while retaining MS/MS spectra similarity for relationships.
- But in addition, relationships in networks can be established between *at least* two objects. So where a network, represented mathematically as a graph, is limited to recording relationships between pairs of objects, a *hypernetwork*, mathematically represented as a multidimensional, multiway *hypergraph*, is not so restricted.

The move from networks (graphs) to hypernetworks (hypergraphs) is a burgeoning movement in data science, and hypergraph methods have been applied to address problems in a variety of fields, including complex physical systems,^5,6^ epidemiology^7,8^ and computational transcriptomics.^9^

Accordingly, in this work we introduce “molecular hypernetworks” (MHNs) as an extension of MNs (including FBMNs) to support the representation of multiway, multidimensional edges describing connectivity among MS features, and provide some initial indications of their utility in metabolomics. Our MHN approach extends MNs by: (1) extending graph edges on molecular pairs in MNs to multi-way hyperedges among groups of molecules in MHNs; and (2) considering combinations of MS dimensions, including drift time, retention time, precursor *m/z*, and MS/MS to analyze multi-dimensional similarity.

The most obvious utility of MHNs relative to MNs is their ability to augment exploratory data analysis activities through consequential improvements to visual interpretability. MHNs visually simplify highly connected regions. In a MHN groups of nodes that are fully connected are grouped by a hyperedge, reducing the number of lines in the graph. This decrease in visual noise ultimately eases the interpretation of the underlying data. We will also note several opportunities to bring network science analytical methods, algorithms, and measures to bear on such representations, including hypergraph methods such as path and component analyses where length and “width”, or the amount of overlap among the hyperedges of a hyperpath,^10^ can offer analytical insights (see “Networks and Hypernetworks” section below).

Finally, one of the most promising aspect of MN representations is their ability assist with “annotation”, that is, the association of a particular molecular identify to a certain hyphenated MS data feature. More specifically is the potential to assist with creating new annotations by transferring known annotations of features to features that are very similar, but have not yet been annotated. In this context, we can interpret feature similarity as distance in the MN or MHN. But in an MHN, two molecules are no longer only either connected (similar) or not, as they are in an MN, but they can participate in multiple multi-way similarities (hyperedges). Annotations can also be propagated using the MHN structure, but the existence of multiple multi-way similarities among pairs of molecules provides added confidence for annotation propagation. In this way our work aims ultimately to increase confidence in annotation propagation by extending MNs to MHNs.

In this paper we first introduce the essential mathematical concepts of graphs and hyper-graphs on a formal basis, including illustration of simple examples. This includes the “clique reconstruction” method of creating a hypergraph representation of a traditional network, in which each maximal *k*-clique (set of *k* vertices, all pairwise connected) is transformed into a hyperedge of size *k*, thereby producing a hypergraph. We then define MHNs in the context of both GNPS networks and hyphenated MS data generally. We first introduce a specific MHN we built in the context of one particular relational view of a reference data set. We then describe MHNs derived as clique reconstructions of MNs and FBMNs as used in three prominent applications of FBMNs as they relate to samples involving human plasma, the gut microbiome, and plant extracts for the purpose of therapeutic discovery.^1,2,11^ We will demonstrate the ability of the MHNs to elucidate connections between groups of features present in the data, to aid in both discrimination and annotation.

## Methods

### Networks and Hypernetworks

The structures and methods we introduce rely on the mathematical notions of graphs and hypergraphs (or “networks” and “hypernetworks”). Before we define *molecular* networks and hypernetworks, in the following subsections we provide a brief and basic introduction to the mathematical structures and notation here. Conceptually, graphs are used to model pairwise relationships among a collection of entities and are commonly used to interrogate data from networks such as social and collaboration networks (“friend” or “following” relationships), transportation (can one transit from point A to point B directly), infrastructure, and protein interactions, among many others.

Stated generally, a graph starts with a finite set of *vertices*, *V*, also referred to as *nodes*, that represent the entities of interest. Then, if there is a pairwise relationship in the data between two vertices, *v* and *u*, we say that there is an *edge e* = {*v, u*} as an unordered pair of vertices, and a member of an *edge set E*. Visually, we can draw a graph by representing vertices as points on a plane and edges as line segments connecting the points. Having a graph structure and visualization can help make sense of complex data. Being able to see that the vertices are all similarly connected, or that there is one vertex that is highly connected and many that have very few connections, gives insight to global properties of the data. And we can discover new connections or relationships not apparent from the edges themselves when there are indirect connections between two vertices, say *v*_0_ and *v_n_*, as a path *v*_0_*, v*_1_*, . . ., v_n_*, when there are edges {*v*_0_*, v*_1_}, {*v*_1_*, v*_2_}, . . ., {*v_n−_*_1_*, v_n_*} in the graph.

A *hypergraph* then extends the concept of a graph to allow for relationships to be multi-way, not just pairwise. Again we start with a set of vertices, *V*, to represent our entities. But in a hypergraph we consider relationships among any subset *e* = {*v*_1_*, v*_2_*, . . ., v_k_*} ⊆ *V* of the vertices. Note that in this way, our previous graph edges *e* = {*v, u*} ⊆ *V* are also hyperedges, just having size *k* = 2. In this way, we can see that all networks are, in fact, hypernetworks, just those where all relationships are restricted to being pairwise.

But in many cases such a restriction to pairwise relationships results in information loss. This can be easily illustrated in a collaboration setting. Consider a paper authored by three people: Alice, Bob, and Cliff. This certainly implies that Alice and Bob have worked together, Bob and Cliff have worked together, and Alice and Cliff have also worked together. A graph structure with edges {*A, B*}, {*B, C*}, {*A, C*} would be used to model this scenario, as shown in Figure 1(a), where each author is a node, between which a line segment (edge) is placed when two of the authors have worked together. Note that this diagram is equivalent to that of Figure 1(b), where now each connection between a pair of vertices in a graph edge is represented as a group of two vertices, mathematically, a subset of two vertices. However, the graph does not capture the fact that all three people have simultaneously worked together. Indeed, the same graph would capture the case where three papers were written, one with each pair of authors. And in that setting we do not know if the three of them have worked together as a group, only that the pairs have worked together. The authors’ relationship is actually best represented as Figure 1(c), where now there’s a single set of three vertices, indicating the interaction of all three authors.

**Figure 1:**
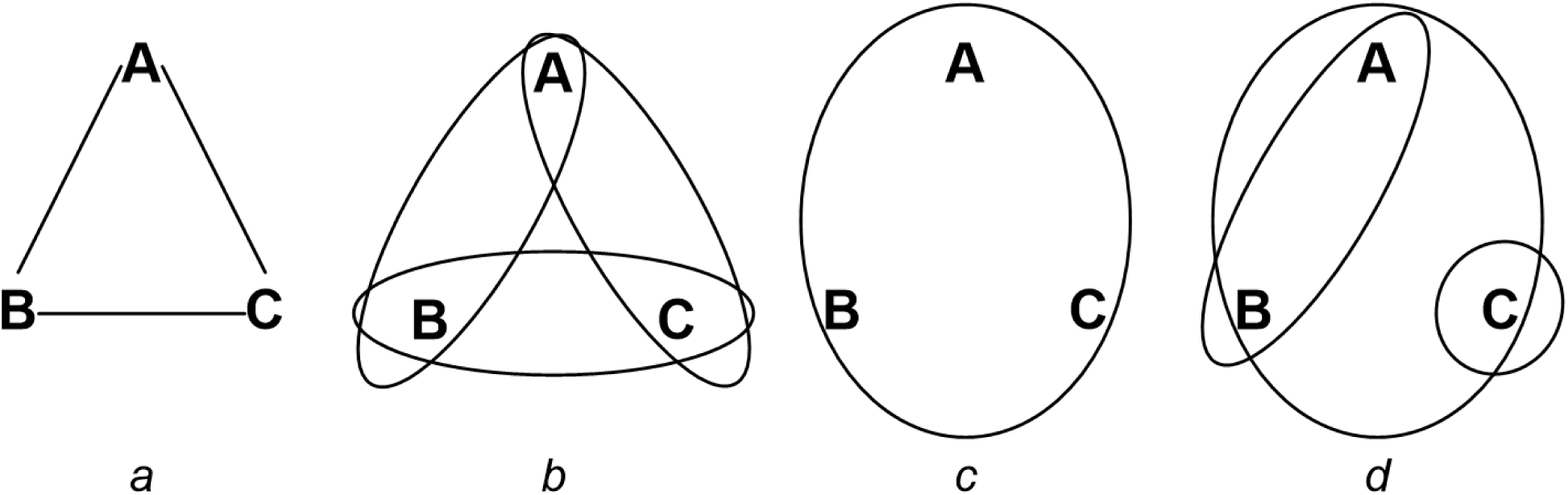
Some example networks and hypernetworks among three elements, vertices *V* = {*A, B, C*}. (a) A graph or network. (b) The same graph shown as a 2-uniform hyper-graph with hyperedge set *ɛ* = {{*A, B*}, {*B, C*}, {*A, C*}}. (c) A proper hypergraph with a single three-way hyperedge, so that *ɛ* = {{*A, B, C*}}. (d) A general hypergraph with three hyperedges combining a proper hyperedge *e*_1_ = {*A, B, C*}, a “graph edge” *e*_2_ = {*A, B*}, and a singleton hyperedge *e*_3_ = {*C*}.

We now introduce these concepts formally. A graph is a structure ⟨*V, E*⟩ with 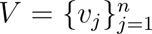 a set of vertices, and 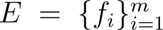 a set of edges, with each edge *f_i_* = {*v, u*}*, f_i_* ∈ *E* a pair of vertices *v, u* ∈ *V* . https://www.overleaf.com/project/63acc9f6070b8e7157ab5da1 Correspondingly, a hypergraph is a structure ⟨*V, ɛ*⟩, where now 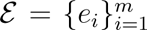 is a family of hyperedges, with each hyperedge a subset *e_i_* ⊆ *V* . Where graph edges involve only two vertices, hyperedges can come in different sizes, |*e_i_*|, possibly ranging from the singleton {*v*} ⊆ *V* (distinct from the element *v* ∈ *V*) to the entire vertex set, so that there is a hyperedge *e_i_* = *V* . It follows that a hyperedge *e_i_* = {*v*_1_*, v*_2_} where |*e_i_*| = 2 is the same as a graph edge, and therefore all graphs are hypergraphs, specifically identified as being “2-uniform”. Figure 1(d) shows a general hypergraph with edges of sizes 1 (a “singleton”), 2 (a “graph edge”), and 3 (a “proper hyperedge”). Figure 1(a) and Figure 1(b) are equivalent, showing the graph and hypergraph representation of the same network, respectively.

Additional mathematical concepts^10^ that are important for this paper include the **degree** of a vertex, which is the number of hyperedges in which it is a member. While vertex degree in a hypergraph is a strict correlate of vertex degree in graphs, hypergraphs are distinguished in that not only can hyperedges have different sizes, but also two hyperedges can intersect in more than one vertex. If two hyperedges *e*_1_*, e*_2_ overlap in *s* vertices, so that |*e*_1_ ∩ *e*_2_| = *s*, then we say that they are *s***-incident**. Graphs are often characterized by their **components**, or collections of vertices such that one can travel from any vertex to any other in the collection via a path of edges between other vertices also in that collection. In hypergraphs, we can additionally require that paths traveled are through edges that have at least *s*-incidence for some *s* ≥ 1. If there is a sequence of edges *e*_1_*, e*_2_*, . . ., e_n_* such that |*e_i_* ∩ *e_i_*_+1_| ≥ *s* for all 1 ≤ *i* ≤ *n* − 1 then *e*_1_ and *e_n_* are said to be *s***-connected**. Then an *s***-component** is a set of vertices implied by edges that are all *s*-connected. As such, 1-connectivity is the equivalent of “regular” connectivity in graphs. So where the left side of Figure 2 shows a single graph component (1-component), in the center the set of vertices {*A, B, C, D, E*} compose the edges *e*_1_*, e*_2_, which are connected by two vertices *B, C*, and thus make a 2-component. Identifying *s*-components is important to discover those portions of the data set that have more dense interactions, indicated by higher *s* overlaps, and is a property unique to MHNs relative to MNs.

**Figure 2:**
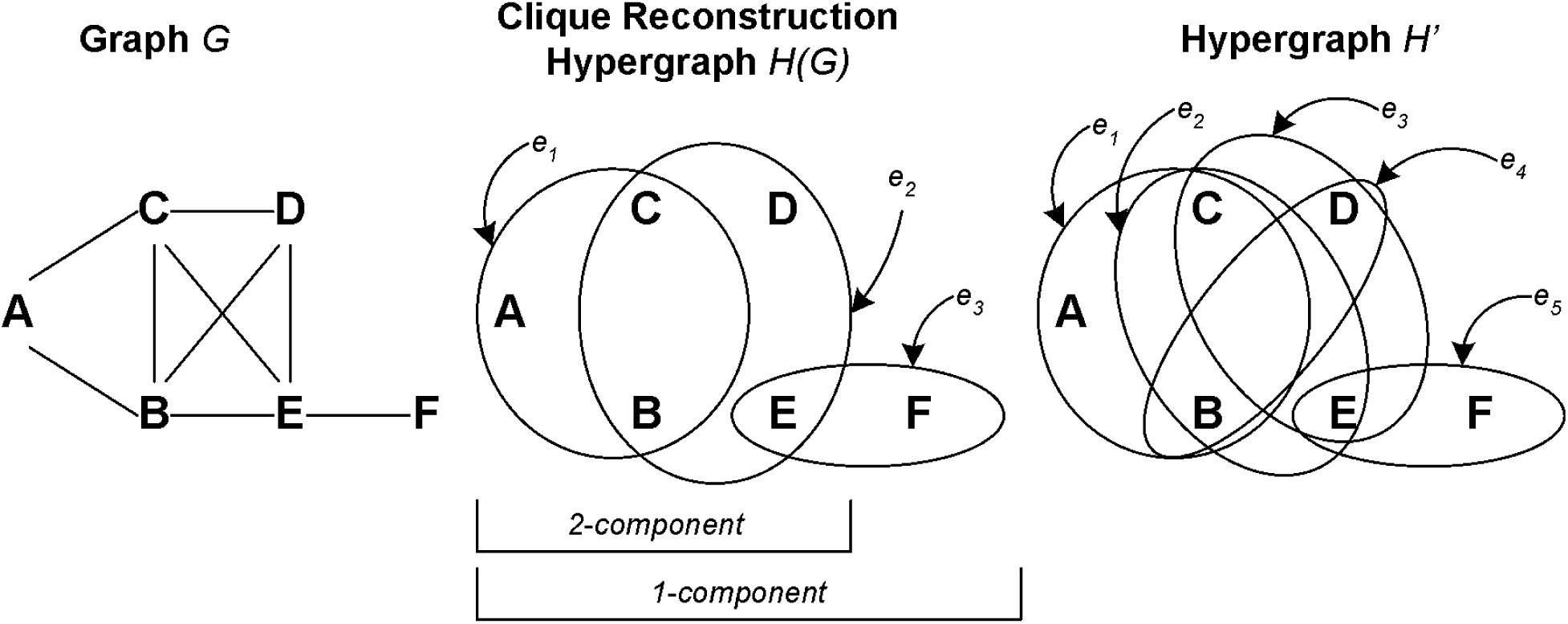
(Left) An example network (graph) *G* containing 9 edges in a single component (1-component). (Center) The hypergraph *H*(*G*) which is the clique reconstruction of *G*. Each of the three 3 maximal cliques of *G* is replaced by a hyperedge *e* ∈ *ɛ*, which now has a 2-component *e*_1_ ∪ *e*_2_ and a 1-component *e*_1_ ∪ *e*_2_ ∪ *e*_3_. *G* is the underlying graph of *H*(*G*), and note that while the graph degree of *C* in *G* is 4, the hypergraph degree of *C* in *H* is only 2. (Right) An example of a different hypergraph *H^′^* with 5 hyperedges. *H^′^* has the same underlying graph *G* as does *H*(*G*), and *H* is the minimal such hypergraph which does.

### Clique Reconstruction Hypergraphs

This paper uses the **clique reconstruction** method to build a hypergraph *H*(*G*) from a graph *G*, which works as follows. For any given graph *G* = ⟨*V, E*⟩, we can identify a **clique** of *G* as a set of vertices *U* ⊆ *V* that induces a completely connected subgraph, in that each vertex pair {*v, u*} ⊆ *U* is an edge in *E*. A **maximal clique** of *G* is then a clique *U* ⊆ *V* such that there is no other clique *U ^′^* of *G* in which *U* is contained. Given an input graph *G*, then its clique reconstruction is a hypergraph *H*(*G*) = ⟨*V*, *ɛ*⟩ where *U* is an edge in *ɛ* if and only if *U* is a clique in G. In other words, every clique in *G* becomes an edge in *H*(*G*) and every edge in *H*(*G*) comes from a clique in *G*.

In the other direction, for any given hypergraph *H* = ⟨*V*, *ɛ*⟩, we can construct its **underlying graph** as a graph *G* = ⟨*V, E*⟩ on the same set of vertices where a pair {*v, u*} of vertices is an edge *f* ∈ *E* if and only if there is a hyperedge *e* ∈ *ɛ* such that *f* = {*v, u*} ⊆ *e*. In other words, the underlying graph *G* is the graph on the same set of vertices *V* that is “implied” by the hypergraph *H*, in that all possible pairwise relationships present in *H* are included.

The clique reconstruction *H*(*G*) of a graph *G* is the minimal hypergraph that has *G* as its underlying graph. It follows that the underlying graph of the clique reconstruction *H*(*G*) of a graph *G* is equal to *G*. But, there can be other hypergraphs *H^′^* ≠ *H* on *V* that also have *G* as their underlying graph. And it is *not* generally the case that the clique reconstruction *H*(*G*) of the underlying graph *G* of a given hypergraph *H* is equal to the original hypergraph *H*. An example is shown in Figure 2, where both of the hypergraphs in the middle and right have the graph on the left as their underlying graph, whereas the clique reconstruction of that graph is only the hypergraph in the center.

Central to this method is the recognition that the clique reconstruction *H*(*G*) captures all of the information in the underlying graph *G*, but does so in a much simplified format. Considering *H*(*G*) shown in the center of Figure 2, its base graph *G* on the left is recovered simply by recognizing that all pairs of vertices in each hyperedge are connected in *G*. But *H*(*G*) has only three hyperedges, compared to the nine graph edges in *G*. And additional information is also available in the hypergraph *H*(*G*). Most importantly is that each of the hyperedges (maximal cliques) *e*_1_*, e*_2_ and *e*_3_ should be considered as a distinct entity, representing all the mutual interactions of the vertices. And beyond that, the *strength* of any relationship between these new entities is also recognized: in the figure, the three vertices *A, B, C* making up the hyperedge *e*_1_ share two vertices with *e*_2_, while *e*_2_ shares only one vertex with *e*_3_.

### Molecular Networks and Hypernetworks

In the original GNPS application, ^3^ MNs are produced by representing measured MS/MS spectra as vectors 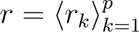 of MS/MS fragment intensities, where now *p* is the number of fragments for a precursor molecule. Binning intensities according to *m/z* results in vectors of equal length *p* for purposes of comparison. Between any two measured spectra *r, r^′^*, a similarity *S*(*r, r^′^*) is calculated as the cosine similarity

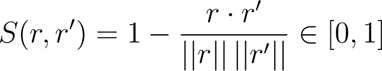

of their MS/MS fragmentation spectra, where · is the vector dot product and ||*r*|| is the Euclidean norm. *S*(*r, r^′^*) = 1 when the spectra are identical, and 0 when there is no similarity. GNPS further uses a *modified* cosine similarity, where the mass difference between precursor masses is taken into account.^12^

To construct a MN, the set of measured MS/MS spectra are cast as the set of vertices, 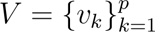, so that each vertex, *v_k_*, represents a spectrum, *r_k_*, of the same index *k*. Edges are created between vertices in the MN if the spectra that they represent are similar enough.

Spectra that are very far apart in the modified cosine similarity are not considered good candidates to represent structurally similar molecules, and thus are not included as an edge in the MN, whereas those that are close are. For any particular data set and study, it needs to be determined what is “close enough” for two spectra to be seen as linked in the network, and that is set as a tolerance or minimum weight *w* ∈ (0, 1]. Retaining pairs of spectra for which *S*(*r_i_, r_j_*) ≥ *w* results in a graph structure *G* = ⟨*V, E*⟩, where *E* = {{*v_i_, v_j_*} : *S*(*r_i_, r_j_*) ≥ *w*} is a set of graph edges.

In GNPS,^1^ Feature-Based Molecular Networks (FBMNs) are constructed by a similar method, but the vertices *v* ∈ *V* represent pairs of MS/MS spectra consolidated over joint LC-MS retention times.

### Data Preparation

We now describe the two kinds of MHNs considered in this paper: an MHN developed *de novo* from internal standards data, and MHNs developed as clique reconstructions of GNPS FBMNs.

### Internal Standards

To explore construction of MHNs using separation dimensions (e.g. ion mobility drift time, liquid chromatography retention time), we analyzed samples containing stable isotope labeled internal standards added to plasma and liver samples. Standards were obtained from a mix of NIH standards representing nine metabolite classes and the QreSS kit from Cambridge Isotope Laboratories (MA, USA). The samples were chromatographically separated using hydrophilic interaction liquid chromatography (HILIC) and reversed-phase liquid chromatography (RPLC) chemistries in separate injections. The LC system was coupled to an Agilent 6560 ion mobility quadrupole time of flight mass spectrometer (Agilent Technologies, CA, USA), and the standards were analyzed in both positive and negative ionization mode.

The MS/MS data were acquired at collision energies of 10, 20, and 40eV using an all-ions fragmentation approach with frames alternating between high and low fragmentation using a mass range of 50-1500 *m/z*.

### Published Feature Based Molecular Networks

Next we were interested in comparing MHNs against prominent MNs and FBMNs as used in the literature. We reanalyzed three networks previously analyzed by feature-based molecular networking^1^ using MS/MS spectra similarity, comparing the resulting MHN to the original MN.

#### Plasma

The first network was derived from analysis of human plasma (MassIVE ID MSV00008263) collected on an ultra-high performance liquid chromatography device (Vanquish, Thermo Fisher Scientific, MA, USA) coupled to an Orbitrap mass spectrometer (Q Exactive, Thermo Fisher Scientific, MA, USA). The GNPS FBMN was constructed from 20,853 positive-mode features, related by modified cosine similarity of respective MS/MS spectra. Network edges were filtered to have a cosine score above *w >* 0.65 and more than 3 matched peaks. A subcomponent of the FBMN containing Ethylenediaminetetraacetic acid (EDTA) was considered for comparison to the MHN-based approach.

#### American Gut Project

The second network was constructed from the analysis of fecal samples from the American Gut Project (MassIVE ID MSV000080179) using a reversed-phase high-performance liquid chromatography device (Dionex UltiMate 3000, Thermo Fisher Scientific, MA, USA) coupled to a quadrupole time of flight mass spectrometer (Impact HD, Bruker, MA, USA). ^11^ The GNPS FBMN was constructed from 194,528 positive-mode features, related by modified cosine similarity of respective MS/MS spectra. Network edges were filtered to have a cosine score also above *w >* 0.65 and more than 5 matched peaks. A subcomponent of the FBMN containing several N-acyl amide isomers was used for comparison to the MHN-based approach.

#### Euphorbia Dendroides

Finally, the third network was created from analysis of 14 *Euphorbia dendroides* plant sample extracts (MassIVE ID MSV000079856) using an HPLC Ultimate 3000 system (Dionex, Voisins-le-Bretonneux, France) coupled to a hybrid linear ion trap/Orbitrap mass spectrometer (LTQ-XL Orbitrap, ThermoFisher Scientific, Les Ulis, France).^2^ The GNPS FBMN was constructed from 7,383 positive-mode features, related by modified cosine similarity of respective MS/MS spectra. Network edges were filtered to have a cosine score above *w >* 0.7 and more than 12 matched peaks. A subcomponent of the FBMN containing 4-deoxyphorbol esters was used for comparison to the MHN-based approach. For each, the FBMN was obtained from GNPS, from which MHNs were constructed by way of clique reconstruction, detailed in the following section.

### Building Molecular Hypernetworks

All MHNs in this study were represented in PNNL’s HyperNetX (HNX) hypergraph modeling platform^†^. For any hypergraph *H*, HNX supports identification of all *s*-components (1 ≤ *s* ≤ 10), which can be visualized in HNX’es hypergraph visualization “widget” tool^‡^, along with relevant statistics (see Figure 4 below).

To build the clique reconstructions of the three studies’ FBMNs, each network *G* was initially loaded with GNPSDataPackage^§^ and processed with the NetworkX graph package^¶^ to extract all maximal cliques *U* ⊆ *V* . These were then established as hyperedges by clique reconstruction in each *H*(*G*) in HNX.

We have produced code and an interactive Jupyter notebook for converting GNPS FBMNs to MHNs via clique reconstruction. The exploratory data analysis workflow provides utility functions for filtering data (e.g., by parent mass, consensus retention time, compound name, and adduct) and searching for hypergraph *s*-components containing nodes with the filtered values. In some cases HNX was extended to meet the needs of this work and, additionally, notebooks specific to metabolomics and GNPS graphs were developed.

To complement the interactive widget, we provide an extra utility for interfacing with the GNPS Browser Network Visualizer, to view the original MN source, and Metabolomics Spectrum Resolver^‖^, to view the MS/MS spectra associated with any node.

## Results

### Internal Standards

As introduced above, there is great flexibility to represent complex, multi-dimensional relational data in different “views” in an MN or MHN. To both show this flexibility in MHNs and introduce their basic properties, we used our internal standards data to generate a *de novo* MHN that cannot be generated as the clique reconstruction of a GNPS MN. ^13^ A portion is shown in Figure 3, where vertices *v* ∈ *V* are distinct precursor *m/z* values, and hyperedges *e* ∈ *ɛ* surround groups of *m/z* values with the same LC-IMS (drift time, retention time) pairs. Nodes are colored by *m/z*, and sized by intensity aggregated over the three feature dimensions. Edges are colored by retention time.

**Figure 3:**
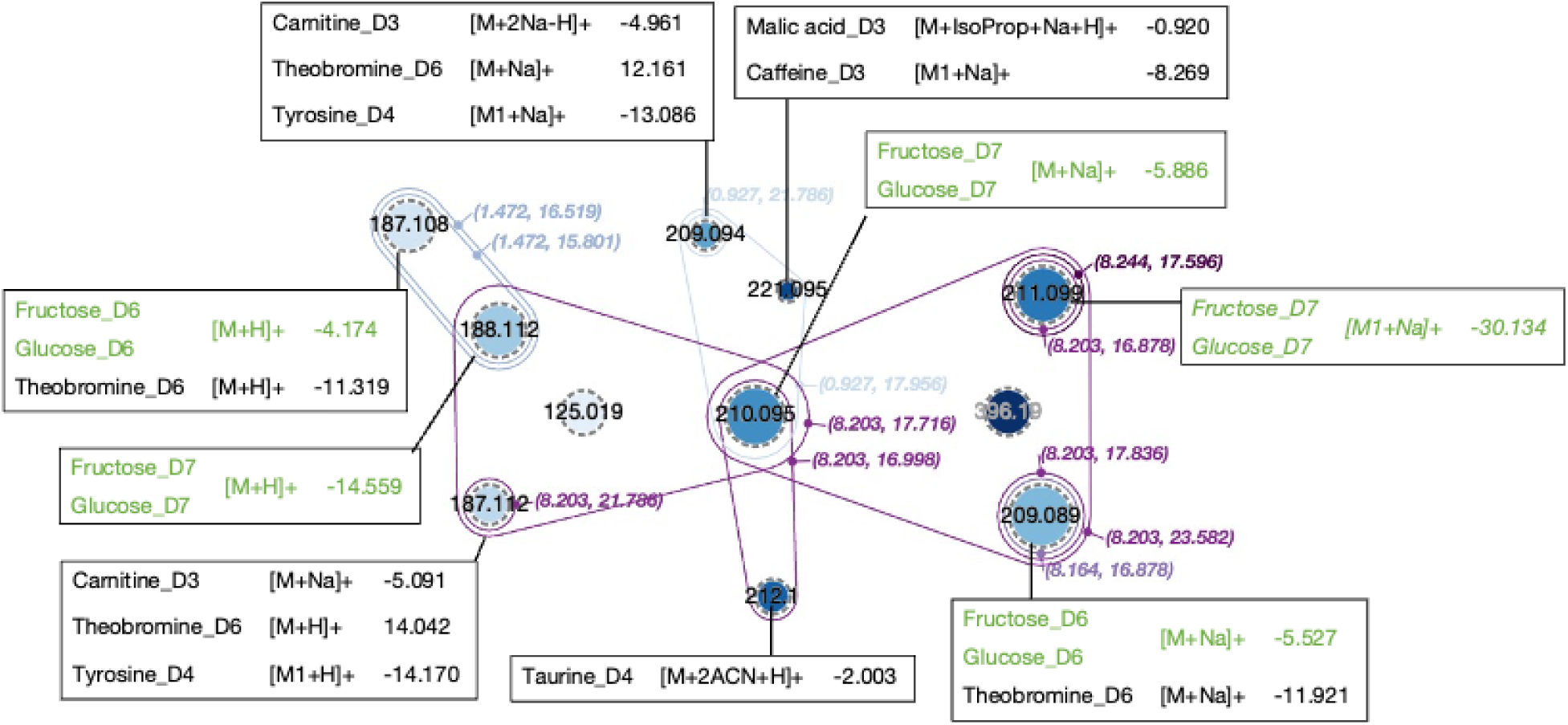
A component of a molecular hypergraph representation of observations of internal standards data. Here nodes indicate distinct precursor *m/z* values. Nodes are grouped into hyperedges when they have the same drift and retention times, as indicated by the pairs (*d, r*) annotating each hyperedge. Candidate annotations are shown. The value after the candidate annotation represents the expected number of deuterium substitutions. For example, “Fructose_D7” denotes a fructose molecule with seven deuterium substitutions. Any putatively observed isotopologues are designated relative to the monoisotopic mass, e.g. [M1+H]+ has one additional neutron compared to [M+H]+. Any putative annotation associated with fructose or glucose has green text (see discussion in text).

Candidate annotations were assigned using BEAMSpy^∗∗^, to nodes that were within 15ppm of the mass of a molecule known to be present within the sample. During this step we only considered putative annotations of monoisotopic mass (M) and single-isotope isotopologues (M1). The only exception was the annotation assigned to the feature with mass 211.099, which has a ppm error of 30. A list of adducts that were considered and molecules known to be present in the sample can be found in the Supplemental Information (ESI-MS-adductsM0- 6.csv, NIH_compounds.csv).

The resulting *de novo* MHN gives several lines of evidence for the presence of deuterated fructose and/or glucose (Figure 3). Five different nodes were putatively annotated as being five different adducts of deuterated fructose and/or glucose. Of these five nodes, three of them are annotated as three different forms of sodiated sugars. Specifically, the nodes with *m/z* 210.095 and 211.099 are annotated as the sodiated form and its M1 isotopic peak. The third node (*m/z*=209.089) could be explained by sodiated sugar that underwent deuterium hydrogen backexchange, which has been previously observed in proteomics data. ^14^ Based off these annotations, we would expect all three forms to have the same retention and drift time. This expectation is shown by the MHN in that a large triangular hyperedge on the right, with retention time of 8.203 minutes and drift time of 23.582 milliseconds, connects all three nodes. In addition, the remaining two nodes (*m/z*=188.112 and *m/z*=187.108) were annotated as two forms of the protonated adduct are connected by two different hyperedges. This meets our expectation that molecules that undergo hydrogen deuterium exchange will retain the same retention and drift time.

Finally, a relatively abundant feature with *m/z* 396.190 was not annotated from the known contents of the internal standards mixture. However, this compound shares drift and retention times with fructose-D7 and glucose-D7 isotopologues, signaling the potential to infer annotation by propagation. For example, a sodiated dimer of fructose/glucose-(D6/D7), by way of deuterium-hydrogen back exchange, yields an *m/z* congruent with that of the observed feature; that is, within 8 ppm *m/z* error.

### Example of a Clique Reconstruction Molecular Hypernetwork in HyperNetX

A MN of Euphodendroidin connectivity generated from the Euphorbia Dendroides data using Cytoscape^††^ is shown in Figure 4A. The 1-component of the MHN built from the corresponding clique reconstruction is shown in Figure 4B, generated by the HyperNetX visualization widget. The HyperNetX tool provides both a flexible, interactive environment to position nodes and hyperedges as well as the ability to display annotations, hover to expose all feature dimensions, and panels for overall statistics including degrees, edge sizes, and related attributes.

**Figure 4:**
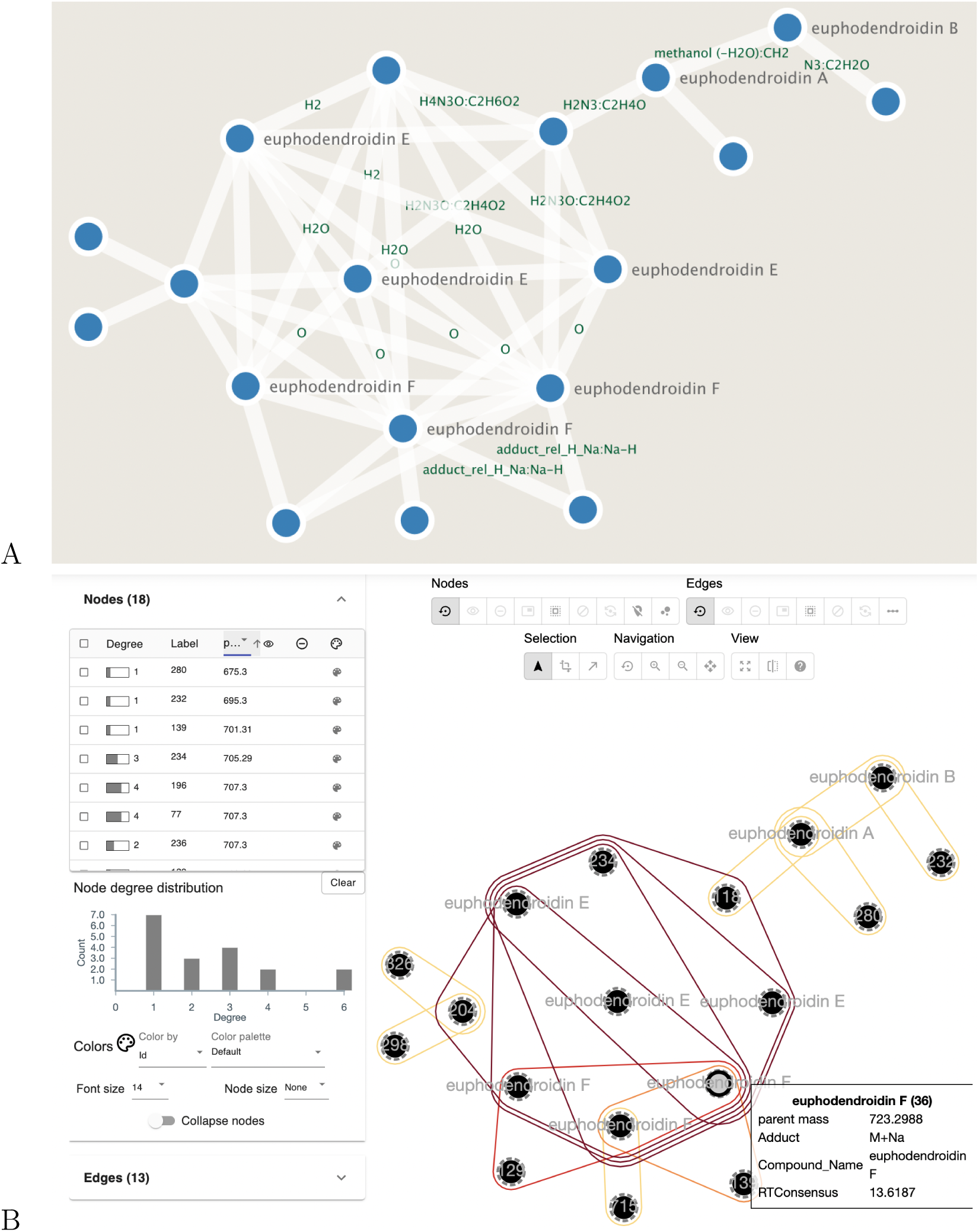
**Euphodendroidin component from the molecular network and hypernetwork for *Euphorbia dendroides* plant samples**^2^ A) MN component in the Cytoscape^15^ interface depicting connectivity of Euphodendroidin . B) Corresponding MHN 1-component in the HyperNetXWidget interface. The left pane contains hypergraph statistics including vertex degree and hyperedge size distributions, while the right pane holds an interactive visualization environment for manipulating the MHN. Additional information is displayed when hovering over a node, as shown.

Note the tremendous advantages for exploration of these data available in this MHN representation in HNX as revealed in Figure 4. The branches of the MN on the left and right of (A) are clearly recreated in the MHN in (B). While the central region of the graph (A) is a much more complicated region, showing great connectivity, it is not complete connectivity. In fact, the maximal clique structure of that central region is revealed in the hyperedges in (B). Indeed, in fact, that region consists of five sub-neighborhoods which are completely connected within them, but not between them. The highlighted node “euphodendriodin F (36)” has maximal degree at 5, indicating that it is included in all five of these regions, but it is the only one so. None of these insights are available in the MN shown in (A).

The overall HyperNetX Python environment additionally provides a wide range of hyper-graph analytical methods, including *s*-components but also extending to a host of hypergraph analytics^10^ that can be used to further analyze both individual and groups of components in the context of the structure of the overall hypergraph. Some of the *s*-component results are shown in Figure 5. The left panel shows the cumulative distribution of the sizes of *s*- components, 1 ≤ *s* ≤ 10 in terms of the total proportion of hyperedges in the MHN. Here we have excluded singleton components consisting of a single edge (of any size) disconnected from others. We can see that such singletons are the bulk (100% − 40% = 60%) of the edges, and also that there are just a few components with high *s* (minimum walk “width” connecting any pair of edges). The central panel shows the size distribution of the components, revealing one large 1-component containing more than 50 hyperedges, but many more with fewer. The right panel shows the distribution of the annotations available from GNPS in the graph over the components, indicating the identity of the component shown in Figure 4.

**Figure 5:**
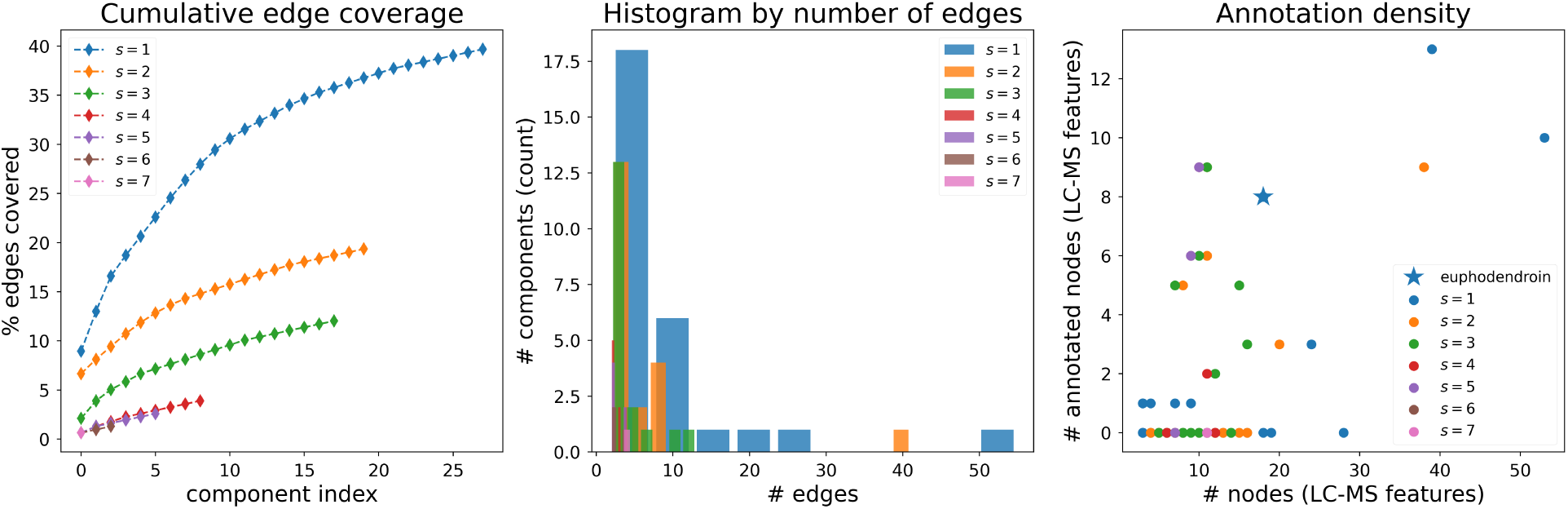
**Non-singleton *s*-component statistics for molecular hypernetwork for *Euphorbia dendroides* plant samples.**^2^ Euphodendroidin component shown in Figure 4 is marked with a star in the annotation density plot (right).

### Clique Reconstruction Molecular Hypernetworks

We now illustrate some properties of molecular hypernetworks by displaying the clique reconstructions of GNPS-derived MNs in Figures 6, 7, and 8. As discussed above, we reanalyzed three datasets originally presented in the paper introducing feature-based molecular networking^1^ by building their clique reconstruction MHNs. As noted above, functionally, this means that groups of nodes that are fully connected, *i.e.,* all pairs of are connected to each other in a MN, are grouped by a hyperedge. We will see that the relationships described by a standard MN are contained within the MHN. And while the overall structure of the MN is recreated by a MHN, it can offer additional clarity in depicting multi-vertex relationships. When displayed, this can replace a large group of individual binary edges in the MN with a single hyperedge polygon in the MHN to display them as a collection of highly similar nodes.

**Figure 6:**
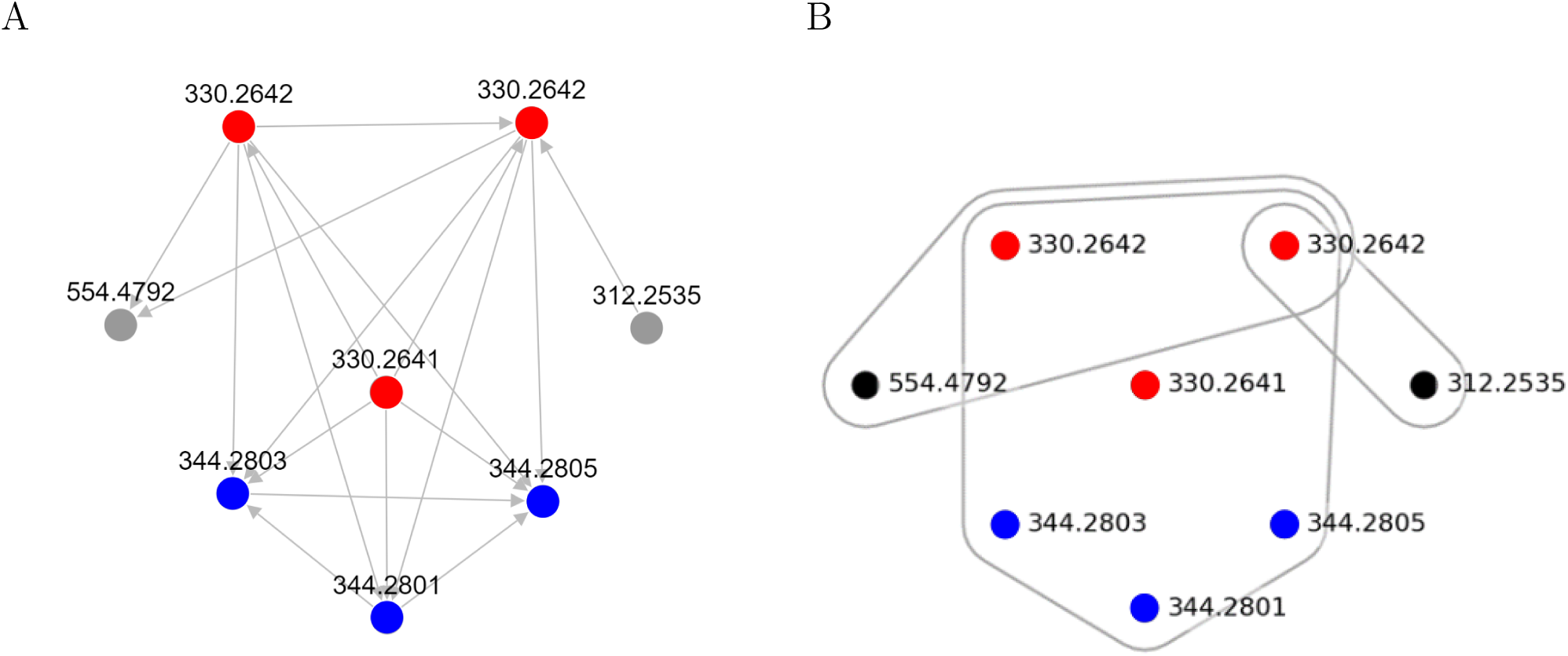
**Comparison of molecular network and molecular hypernetwork using data from the American Gut Project** (A) Molecular network view of N-acyl amides found in gut samples (Figure 2-d in the original publication^1^). Nodes indicate the precursor m/z and edges correspond to relationships with cosine score > 0.65 and number of matched peaks > 5. Colored nodes indicate two groups of isomers: 330.26 m/z in red and 344.28 m/z in blue. (B) Clique reconstruction molecular hypernetwork view of the same data found in (A). Both kinds of networks are able to represent the same information for simple networks like this example, but the hypernetwork is visually simpler. Rather than pairwise relationships represented by multiple edges in (A), a single hyperedge in (B) captured all relationships within the related molecules and directly represents the high spectra similarity among the 2 set of isomers in a simpler way.

An analysis of the network containing N-acyl amides in fecal samples illustrates how the biological insight gained from networks are realized in hypernetworks (Figure 6), and perhaps better so in terms of simplified and clearer visual representations. In this component, two sets of isomers share high spectral similarity. One set of three nodes were previously identified as N-(hydroxyhexadecanoyl)glycine isomers (*m/z*=330.2642) and the other set of three nodes were identified as N-(hydroxyheptadecanoyl)glycine isomers (*m/z*=344.2801).^1^ These two sets of isomers are very similar as one set of compounds has a slightly longer carbon chain, and therefore their MS/MS is expected to be similar. All the similarity values are above the cosine similarity threshold, but there are specific fragments driving the similarities within each set (supplemental Figure 1). Both the MN (Figure 6A) and MHN (Figure 6B) showed the similarity within and between the two sets of isomers. However, rather than 15 edges in a MN, the MHN more clearly showed, via a single hyperedge encompassing all six nodes, that the two sets of isomers all have similar MS/MS. Two other clusters of relationships can be observed in the hypernetwork due to additional shared fragments which are unique to the other hyperedges or due to more similar abundance patterns (supplemen-tal Figure 1). The 554.47 *m/z* ion is related to two isomers due to additional fragments at the low mass range. The 312.25 *m/z* ion, annotated as a putative new derivative, N- (dehydrohexadecanoyl)glycine, has a cleaner MS/MS and is related to a single isomer, due to the more similar abundance pattern at the low mass fragment ions.

Figure 7 depicts an FBMN graph centered on the [M+H]+ ion *m/z* = 293.0978 for the chelator EDTA. In-source fragment ions are observed including masses 247.0926, 160.061, 132.0808, and 114.055. The NIST MS/MS library contains data for this ion, which indicates that 293 gives rise to 247, 247 gives rise to 160, and 160 gives rise to 132 and 114 fragment ions. This MSn experimentally-derived fragmentation tree is partially reflected in the network structure, with 293, 247, and 160 sharing membership in a hyperedge, and the fragment ions 132 and 114 more peripheral in the MHN. In the MN, EDTA (*m/z*=293.0978) is connected to an in-source fragment ion (*m/z*=247.0926) derived from EDTA via a single edge. However, in the MHN, these two ions share membership in four different hyperedges, which we call their “adjacency”.

**Figure 7:**
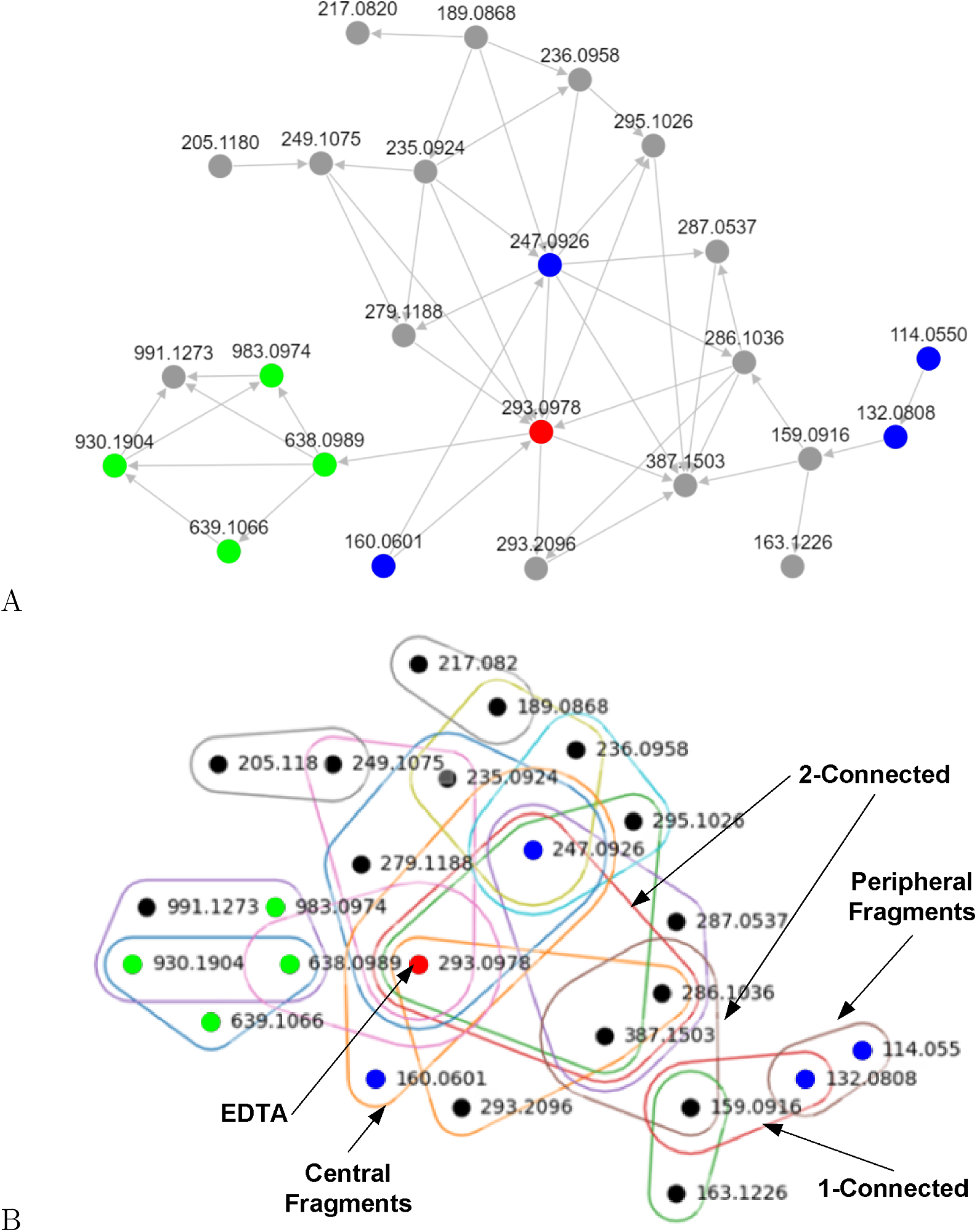
**Comparison of molecular network and molecular hypernetwork for EDTA** (A) A molecular network view of the network around centered around in EDTA that was derived from Figure 2F of *Nothias et al* .^1^ (B) A molecular hypernetwork view of the same data found in (A). Note that edge colors are for visualization purposes only. In addition, note that the red node is EDTA, blue nodes are in-source fragments of EDTA, and green nodes are possible manganese adducts.

Additional network observations are possible as well. For example, in the MN we can observe that the minimal path from the central fragments to the peripheral fragments goes through a path involving the 387 node for the 160 fragment, and 286 for the 247 fragment, and then the 159 node for both. But in the MHN we can see that that chain involves 2-connectivity for the first step, and then 1-connectivity to reach the peripheral fragments. This level of specificity may reveal hidden relationships or putative identities of other measured features.

Further, the left side of the MN and MHN in Figure 7 contains a small sub-network composed of ions 638, 639, 930, 983, and 991. While this was not annotated in the original manuscript, the ions at 638.0989, 639.1066, and 930.1904 can be putatively annotated as manganese adducts [2M0 + Mn - H]+, [2M1 + Mn - H]+, and [3M0 + Mn - 3H]+, respectively. These three adducts are found in a single hyperedge (blue) which partially overlaps with the purple hyperedge containing 638, 930, 983, and 991. While we could not assign a putative annotation to the signal at m/z 991, the signal at 983.0974 is consistent with [3M0 + 2Mn - 3H]+. All putative annotations are consistent at < 20 ppm mass error. The hyperedges of the manganese multimers potentially reflect mechanstic processes of ion formation: the primary [M+H]+ ion is most closely associated with the M0 isotope of the manganese dimer [2M0 + Mn - H]+ and more distal to the doubly adducted trimer [3M0 + 2Mn - 3H]+. This pattern has the potential to reflect ion cluster formation mechanisms, as the 2Mn adduct is unlikely to form without prior formation of the single manganese dimer. While these interpretations will require further validation, they highlight the potential of hypernetworks to improve interpretation of FBMN. Electrospray-ionization based chelation behavior has been validated in other studies, ^16^ lending further support to this interpretation.

To show this visual improvement, we compared a MN and MHN view of a highly connected component that resulted from an analysis *Euphorbia dendroides* plant samples. Specifically, we focused on a highly connected subcomponent of 10 nodes that contained six 4- deoxyphorbol ester isomers (lower right of Figure 8A and Figure 8B). While the MN readily showed that the component is highly connected, it was difficult to determine whether the subcomponent is fully connected (Figure 8A). On the other hand, the MHN easily showed that while these 10 nodes are not fully connected, a subset of eight nodes (blue edge) and seven nodes (brown edge) are fully connected (Figure 8B).

**Figure 8:**
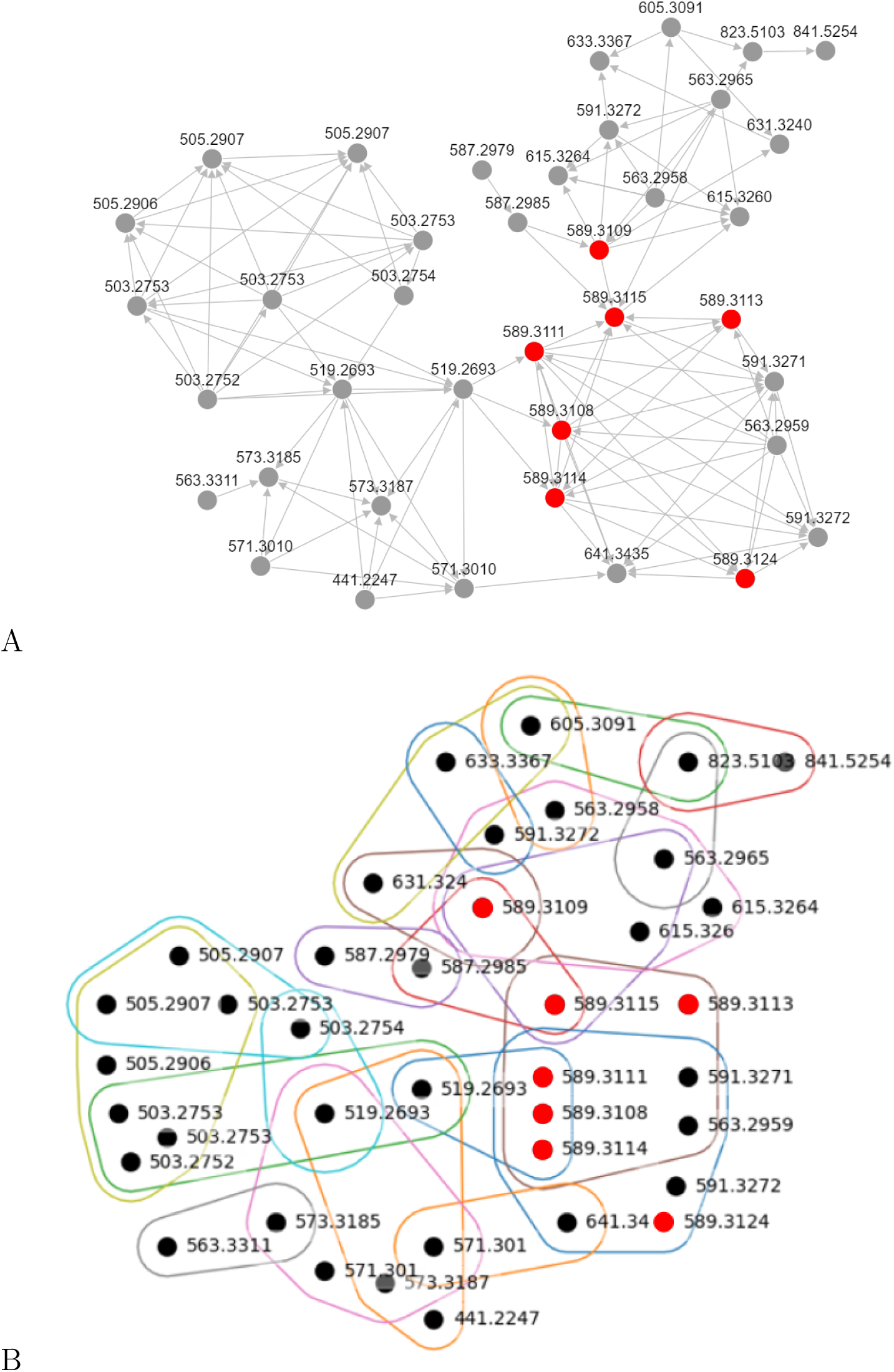
**Comparison of molecular network and molecular hypernetwork for** *Euphorbia dendroides* **plant samples** (A) A molecular network view of 4-deoxyphorbol esters (red nodes) that was derived from Figure 2B of *Nothias et al* .^1^ (B) A molecular hypernet-work view of the data in (A). Note that edge colors are for visualization purposes only. In this example, hypernetworks improve interpretability among highly connected vertices with a high degree of edge occlusion in the MN representation.

## Discussion

In a conventional graph, if there are multiple relationships among a group of nodes, each relationship must be represented by a separate edge, which can lead to a cluttered visual representation. In a hypergraph, a single hyperedge can connect multiple nodes, which simplifies the visual representation by reducing the number of edges. For example, consider the network depicted in Figure 8: the MN representation (Panel A) has a high degree of inter-connectedness among nodes representing 4-deoxyphorbol esters, yet the explicit relationships are difficult to interpret due to edge occlusion. This enables immediate interpretability of the apparent many-way relationships, as well as the degree of relatedness between collections of nodes. More generally, this representation of hierarchical relationships, wherein a node can be a part of multiple hyperedges, is more readily apparent by hypergraph visualization.

Additionally, the connectivity relative to a feature of interest arises naturally from hypernetwork construction. Again consulting Figure 8, the MN representation generated in Nothias et al. truncates the graph relative to the 4-deoxyphorbaol ester isomers. In the MHN representation, hyperedge connectivity resolves to a natural terminus, whereby prospectively relevant features are visualized without stoppage criteria defined by the user.

However, there are challenges that arise from representing connectivity in this multiway fashion, most notably how to transfer annotations often present along binary edges in MN representations to corresponding hyperedges in an MHN derived through clique reconstruction. In the context of GNPS-based structure similiarity MNs, these typically represent differences in *m/z* between connected nodes. The information conveyed by this network adornment has useful implications, such as identifying in-source fragments, changes in functional groups, and other structural motifs, which are currently absent in MHN renderings. We note that the software used to generate the MHNs, HNX, is highly flexible and stewarded by the authors’ institution. As such, a future extension to reveal binary edges and corresponding annotations upon node hover will be developed, achieving a hybrid, best-of-both-worlds solution.

### Conclusion and Future Work

Determining the relatedness of metabolites studied by hyphenated mass spectrometry techniques poses a significant challenge, simplified to some degree by representing detected features as a graph. To advance the interpretation of mutually related features, whether by structural similarity or otherwise, we developed an extension of binary molecular networks to molecular *hyper*networks, enabling multiway, multidimensional connectivity among network vertices. While this effort represents only a proof-of-principle application of hypernetworks to metabolomics, improvements to exploratory data analysis activities, such as visual interpretability, are readily apparent. Additionally, this work serves as a foundational point of departure towards more principled application of network and hypernetwork science. For example, there are a multitude of statistics widely applied in network science to facilitate exploratory data analysis, including path lengths, connectivity analysis, centrality analysis, etc. Congruently, similar metrics exist for hypergraphs, though with differing assumptions and resulting interpretations. For example, the concepts of *s*-connectivity and edge size (the number of vertices contained by a hyperedge) could be used to (i) triage annotation propagation targets, (ii) identify in-source fragments, and (iii) identify chemical background and/or contamination, such as analytes that elute throughout the LC experiment, among others. More advanced topics like hypergraph clustering could be applied to identify groups of related spectra in a new way that respects the varying levels of *s*-connectivity provided when modeling this data as an MHN. Critically, while the literature reviewed for this work mentions the suitability of MNs for analysis by graph/network theory algorithms, such applications have been limited to minimum path length queries.^4,17^ In future works, we intend to further develop MHN functionality to facilitate a “network analysis tool suite”, prospectively easing the process by which data-informed (bio)chemical interpretations are made.

Another forward-looking extension of the MHN representation is to define connectivity relative to multiple representations of the underlying data. While not unique to MHNs relative to MNs, as evidenced by Ernst et al.,^18^ the suitability of multilayer MHN connectivity has yet to be assessed. For example, a structural similarity MHN constructed from MS/MS spectra can be connected to a MHN relating features and adducts to “ion identity”, such as in Schmid et al.^19^ In turn, these networks can additionally connect to a mass difference network, defined by *m/z* deltas among precursor ions, to engender a more complete view of the underlying biochemical relatedness among analytes under study.

In all, MHNs represent a promising tool in the analysis of complex relationships underlying high-dimensional mass spectrometry data.

## Acknowledgements

The authors thank Mingxun Wang, Thomas Metz, and Samuel Purvine for their strong support and assistance during this work. This work was supported by the Pacific Northwest National Laboratory (PNNL) Laboratory Directed Research and Development Program and is a contribution of the m/q Initiative. A portion of the research was performed using resources available through Research Computing at PNNL. PNNL is a multiprogram national laboratory operated by Battelle for the Department of Energy under Contract No. DE-AC05- 76RLO 1830.

## Footnotes

∗ https://gnps.ucsd.edu

† https://pnnl.github.io/HyperNetX

‡ https://github.com/pnnl/hypernetx-widget

§ https://github.com/Wang-Bioinformatics-Lab/GNPSDataPackage

¶ https://networkx.org/

‖ https://metabolomics-usi.ucsd.edu/

∗∗ https://github.com/computational-metabolomics/beamspy

†† https://cytoscape.org/

## References

(1) Nothias, L.-F. et al. Feature-based molecular networking in the GNPS analysis environment. Nature Methods 2020, 17, 905–908.

(2) Nothias, L.-F.; Nothias-Esposito, M.; da Silva, R.; Wang, M.; Protsyuk, I.; Zhang, Z.; Sarvepalli, A.; Leyssen, P.; Touboul, D.; Costa, J.; Paolini, J.; Alexandrov, T.; Litaudon, M.; Dorrestein, P. C. Bioactivity-Based Molecular Networking for the Discovery of Drug Leads in Natural Product Bioassay-Guided Fractionation. Journal of Natural Products 2018, 81, 758–767.

(3) Wang, M.; Carver, J. J.; Phelan, V. V.; Sanchez, L. M.; Garg, N.; Peng, Y.; Nguyen, D. D.; Watrous, J.; Kapono, C. A.; Luzzatto-Knaan, T.;, et al., Sharing and community curation of mass spectrometry data with Global Natural Products Social Molecular Networking. Nature Biotechnology 2016, 34, 828–837.

(4) Amara, A.; Frainay, C.; Jourdan, F.; Naake, T.; Neumann, S.; Novoa-del Toro, E. M.; Salek, R. M.; Salzer, L.; Scharfenberg, S.; Witting, M. Networks and Graphs Discovery in Metabolomics Data Analysis and Interpretation. Frontiers in Molecular Biosciences 2022, 9 .

(5) Battiston, F.; Amico, E.; Barrat, A.; Bianconi, G.; de Arruda, G. F.; Franceschiello, B.; Iacopini, I.; Kéfi, S.; Latora, V.; Moreno, Y. M M. The Physics of Higher-Order Interactions in Complex Systems. Nature Physics 2021, 17, 1093–1098.

(6) Battiston, F.; Cencettib, G.; Iacopini, I.; Latora, V. e. a. Networks Beyond Pair-wise Interactions: Structure and Dynamics. Physics Reports 2020, 874, 1–92, 10.1016/j.physrep.2020.05.004.

(7) Iacopini, I.; Petri, G.; Barrat, A.; Latora, V. Simplicial Models of Social Contagion. Nature Communications 2019, 10, 2485.

(8) Landry, N.; Restrepo, J. G. The Effect of Heterogeneity on Hypergraph Contagion Models. Chaos 2020, 30:*10*, 3117, 10.1063/5.0020034.

(9) Feng, S. et al. Hypergraph Models of Biological Networks to Identify Genes Critical to Pathogenic Viral Response. BMC Bioinformatics. 2021; 10.1186/s12859-021-04197-2.

(10) Aksoy, S. G.; Joslyn, C. A.; Marrero, C. O.; Praggastis, B.; Purvine, E. A. Hyper-network Science via High-Order Hypergraph Walks. EPJ Data Science 2020, 9:*16*, 10.1140/epjds/s13688-020-00231-0.

(11) McDonald, D. et al. American gut: An open platform for citizen science microbiome research. mSystems 2018, 3 .

(12) Watrous, J.; Roach, P.; Alexandrov, T.; Heath, B. S.; Yang, J. Y.; Kersten, R. D.; van der Voort, M.; Pogliano, K.; Gross, H.; Raaijmakers, J. M., et al. Mass spectral molecular networking of living microbial colonies. Proceedings of the National Academy of Sciences 2012, 109, E1743–E1752.

(13) Purvine, E. A.; Bilbao, A.; Broeckling, C.; Colby, S.; Joslyn, C.; Lin, A.; Metz, T.; Shapiro, M. Introducing Molecular Hypernetworks for Exploration of Multi-dimensional Metabolomics Data. 70th American Society for Mass Spectrometry Conf. on Mass Spec-tromety and Allied Topics (ASMS 22), 2022.

(14) Anderson, K. W.; Hudgens, J. W. C for Reduced Back-Exchange, Reduced Carryover, and Improved Dynamic Range for Hydrogen-Deuterium Exchange Mass Spectrometry. J Am Soc Mass Spectrom 2022, 33, 1282–1292.

(15) Shannon, P.; Markiel, A.; Ozier, O.; Baliga, N. S.; Wang, J. T.; Ramage, D.; Amin, N.; Schwikowski, B.; Ideker, T. Cytoscape: a software environment for integrated models of biomolecular interaction networks. Genome research 2003, 13, 2498–2504.

(16) Aron, A. T.; Petras, D.; Schmid, R.; Gauglitz, J. M.; ttel, I.; Antelo, L.; Zhi, H.; Nuccio, S. P.; Saak, C. C.; Malarney, K. P.; Thines, E.; Dutton, R. J.; Aluwihare, L. I.; Raffatellu, M.; Dorrestein, P. C. Native mass spectrometry-based metabolomics identifies metal-binding compounds. Nat Chem 2022, 14, 100–109.

(17) Qin, G.-F.; Zhang, X.; Zhu, F.; Huo, Z.-Q.; Yao, Q.-Q.; Feng, Q.; Liu, Z.; Zhang, G.-M.; Yao, J.-C.; Liang, H.-B. MS/MS-Based Molecular Networking: An Efficient Approach for Natural Products Dereplication. Molecules 2023, 28, 157.

(18) Ernst, M.; Kang, K. B.; Caraballo-Rodríguez, A. M.; Nothias, L.-F.; Wandy, J.; Chen, C.; Wang, M.; Rogers, S.; Medema, M. H.; Dorrestein, P. C., et al. MolNetEnhancer: Enhanced molecular networks by integrating metabolome mining and annotation tools. Metabolites 2019, 9, 144.

(19) Schmid, R.; Petras, D.; Nothias, L.-F.; Wang, M.; Aron, A. T.; Jagels, A.; Tsugawa, H.; Rainer, J.; Garcia-Aloy, M.; Dührkop, K., et al. Ion identity molecular networking for mass spectrometry-based metabolomics in the GNPS environment. Nature communications 2021, 12, 3832.

