## Supplementary figures and images for "Introducing Molecular Hypernetworks for Discovery in Multidimensional Metabolomics Data"

### spectra_312_2535.png

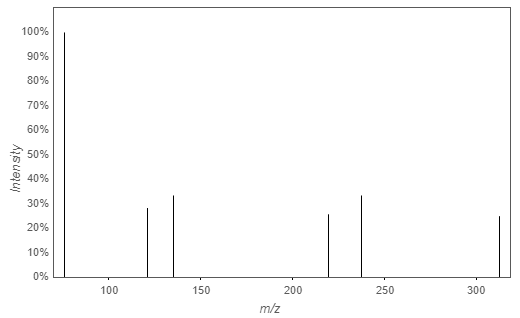

### spectra_330_2641.png

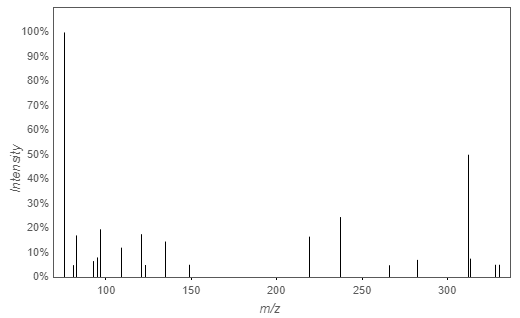

### spectra_330_2642_left.png

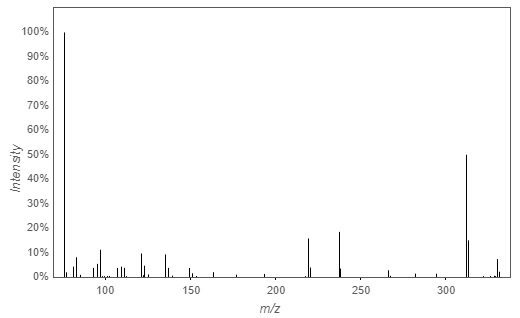

### spectra_330_2642_right.png

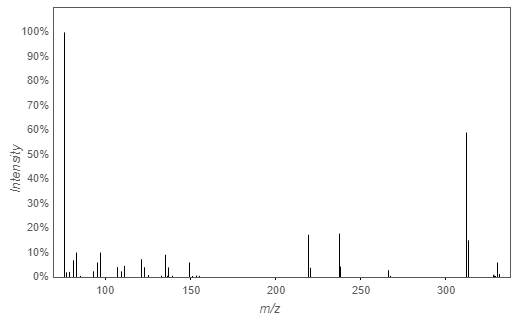

### spectra_344_2801.png

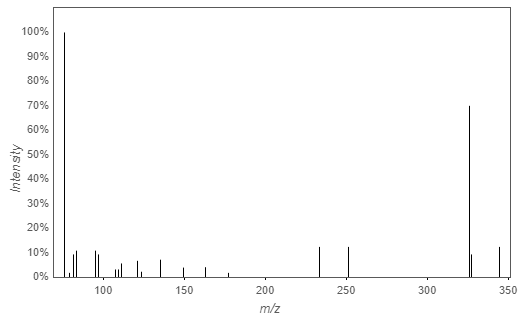

### spectra_344_2803.png

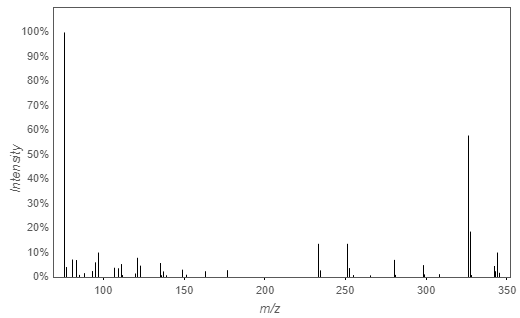

### spectra_344_2805.png

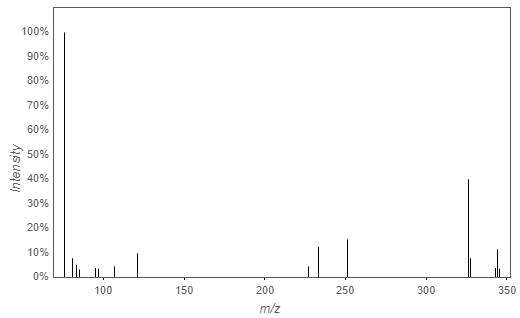

### spectra_554_4792.png

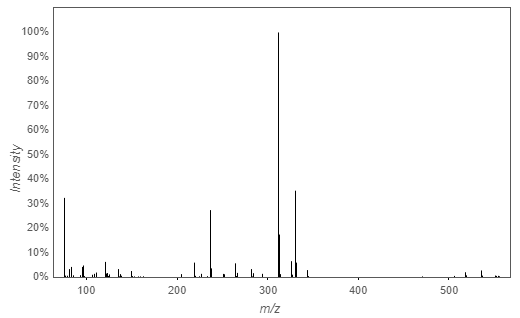
